# Phenological matching drives wheat pest range shift under climate change

**DOI:** 10.1101/614743

**Authors:** Yuqing Wu, Zhongjun Gong, Daniel P. Bebber, Jin Miao, Zhonghua Zhao, Yuying Jiang, Shihe Xiao, Guoyan Zhang, Dazhao Yu, Jichao Fang, Xinmin Lu, Chaoliang Lei, Jianqing Ding, Qiang Wang, Yueli Jiang, Tong Li, Hongmei Lian, Huiling Li, Yun Duan, Jianrong Huang, Donglin Jing, Yunzhuan He, Zhi Zhang, Yunhui Zhang, Julian Chen, Hongbo Qiao, Wenjiang Huang

**Author notes:** These authors contributed equally to this work (Yuqing Wu, Zhongjun Gong, Jin Miao).

## Abstract

Shifting geographical ranges of crop pests and pathogens in response to climate change pose a threat to food security (*1*, *2*). The orange wheat blossom midge (*Sitodiplosis mosellana* Géhin) is responsible for significant yield losses in China (*3*), the world’s largest wheat producer. Here we report that rising temperatures in the North China Plain have resulted in a mean northward range shift of 3.3° (58.8 km per decade) from the 1950s to 2010s, which accelerated to 91.3 km per decade after 1985 when the highly toxic pesticide hexachlorocyclohexane (HCH) was banned (*4*). Phenological matching between wheat midge adult emergence and wheat heading in this new expanded range has resulted in greater damage to wheat production. Around $286.5 million worth of insecticides were applied to around 19 million hectares in an attempt to minimize wheat midge damage to crops between 1985 and 2016. Despite use of these pesticides, wheat midge caused losses of greater than 0.95 million metric tons of grain during this period. Our results demonstrate the potential for indirect negative impacts of climate change on crop production and food security, and constitute the first large scale example of plant pest range shift due to global warming.

---

Latitudinal range shifts of wild species populations are among the strongest biological signals of recent anthropogenic climate change (*5*). Similar range shifts in invasive species or pests and pathogens of agricultural crops and livestock would have significant implications for biodiversity conservation or food production, respectively. While there is some evidence for latitudinal range shifts in line with global warming for crop pests and pathogens in general (*1*), few emerging pest outbreaks have been unequivocally attributed to climate change (*2*). Similarly, while many studies have modelled the effects of climatic change and variability on crop production, the impact of pests and diseases is generally ignored. Given the huge production losses attributable to crop pests and pathogens (*6*), and the economic and environmental burdens resulting from their control, understanding how climate change alters the risk of outbreaks is key to improving global food security in a warming world.

Wheat (*Triticum aestivum*) is the world’s most important crop in terms of production area and second-most important in human diets. Wheat is grown mainly at temperate latitudes in both hemispheres, with China the world’s largest producer. The North China Plain (NCP, Fig. 1, Supplementary Fig. S1), comprising the provinces of Beijing, Tianjin, Hebei, Shandong, Henan, Anhui, Jiangsu, Shanghai, Zhejiang, Jiangxi and Hubei, grew around one tenth of the world’s wheat between 1961 and 2016. Orange Wheat Blossom Midge (hereafter, wheat midge), *Sitodiplosis mosellana* (Géhin) (Diptera: Cecidomyiidae) is a major wheat pest in northern latitudes around the world, including in the NCP (*3*). Wheat midge is univoltine and has a long diapause period during which cocooned larvae overwinter in the soil (*7*). Diapause is broken by winter chilling (*7*), followed by pupation and emergence of adults from the soil in response to warming and other climatic cues (*8*–*10*). In North America and Europe, wheat midge larvae consume the kernels of spring wheat and other Poaceae crops, resulting in serious losses. An outbreak in northeast Saskatchewan, Canada in 1983 caused losses of $30 million across nine municipalities (*11*). Since this time, wheat midge has spread west and south and is now present in many regions of the Great Plains of North America (*12*). Adult midges are able to disperse many hundreds of kilometers on the wind (*13*). In China, one of the first reported outbreaks occurred in Wuxian, Jiangsu province in 1838 (*14*). By the 1950s, wheat midge had spread into the winter wheat production area in the southern region of the NCP between 28° N and 36° N (*15*). Henan province, for example, suffered wheat production losses of 0.352 million tons (7.7 % of production) from an outbreak in 1954 (*15*).

**Figure 1.**
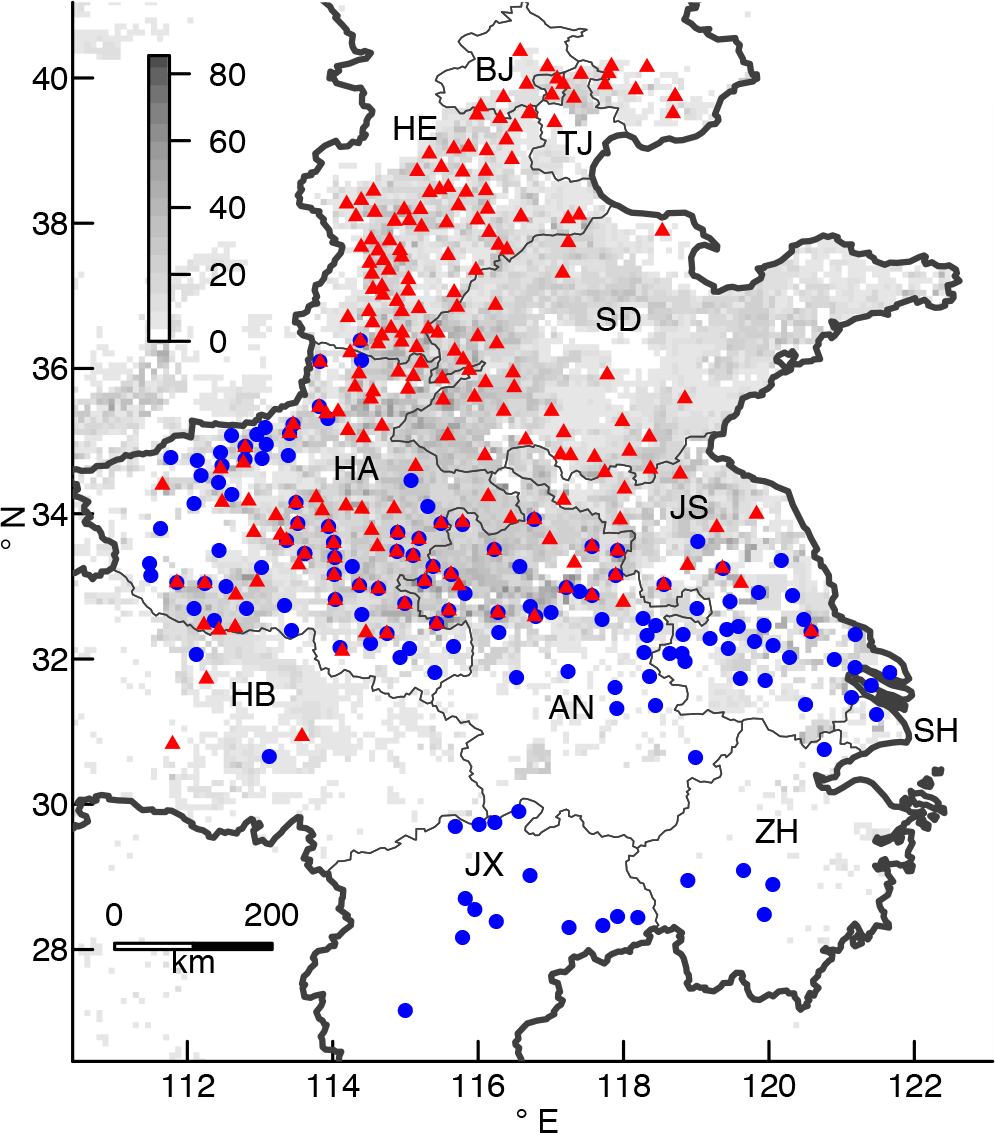
Wheat midge outbreaks in the NCP. Blue circles show outbreaks recorded in the 1950s in the Original Distribution Range (ODR). Red triangles show wheat midge outbreaks from 2010 in the Novel Expanded Range (NER). The NCP is in heavy outline. The grey scale represents the areal wheat planting density in 2005, from MAPSPAM estimates. Provinces in the ODR are Henan (HA), Jiangsu (JS), Hubei (HB), Anhui (AH), Shanghai (SH), Jiangxi (JX) and Zhejiang (ZH). Provinces in the NER are Beijing (BJ), Tianjin (TJ), Hebei (HE) and Shandong (SD). Sea is pale blue.

The history of wheat midge outbreaks in China has been strongly influenced by the pesticide 1,2,3,4,5,6-Hexachlorocyclohexane (HCH), also called benzene hexachloride (BHC) or lindane (*4*). HCH was introduced as a soil treatment in the 1950s, keeping outbreaks in the NCP under control over the period 1958 to 1982 (Fig. 2). Production of HCH in China grew steadily to the mid-1970s, peaking around 250 kt before dropping to zero in 1984 (*4*). The NCP provinces of Jiangsu, Hunan, Henan, Shandong, Hubei, Zhejiang, Jiangxi and Hebei were among the top ten provinces with the highest HCH usage between 1952 and 1984 (*4*). However, serious environmental problems lead to a ban on the application of HCH in 1983 (*4*), resulting in the resurgence in wheat midge and a cumulative loss in wheat production of 2.77 million tons between 1985 and 2016 (Fig. 2). The resurgence of wheat midge was accompanied by a shift in the insect’s geographical range. Given predicted latitudinal range shifts under future climate change for this pest (*16*), and historical range shifts observed for crop pests and pathogens in general (*2*), the altered wheat midge distribution across the NCP over recent decades could be a result of phenological responses to climate change. Here, we investigate the pattern and causes of this range shift, testing the hypotheses that warming temperatures due to climate change allowed the insect to invade the northern regions of the NCP, and that climate change altered phenological matching that is key to successful host exploitation by the pest.

**Figure 2.**
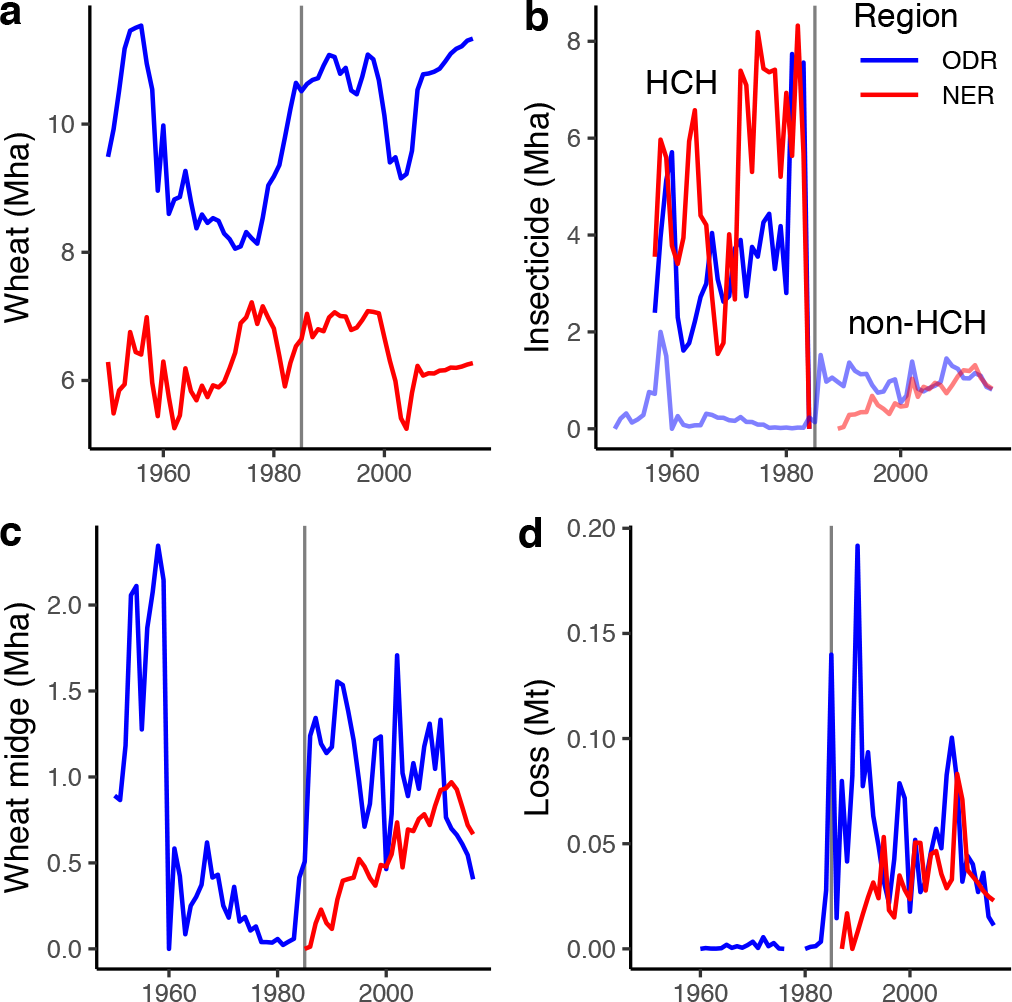
Wheat and the wheat midge in the NCP, 1950 – 2016. a. Wheat production area in the Original Distribution Range (ODR, blue line) and New Expanded Range (NER, red line) provinces (Mha). Grey vertical line at 1985 highlights ban on HCH application. b. Insecticide-treated area (Mha), red and blue show HCH-treated area, pink and pale blue show non-HCH insecticide treated area. c. Wheat-midge affected area (Mha). d. Wheat production loss (Mt).

Wheat midge was restricted to the southern half of the NCP in the 1950s, and the northern half in the 2010s (Fig. 1). Wheat midge damage records from 135 Counties during the period 1950 to 1959 were distributed over the provinces of Henan, Anhui, Jiangsu, Hubei, Jiangxi, Zhejiang and Shanghai across a latitudinal range of 28° N to 36° N. We term this region the Original Distribution Range (ODR). In comparison, 225 reports of wheat midge damage in the NCP were recorded in the 2010s, across eight provinces (Hebei, Beijing, Tianjin, Shandong, Henan, Anhui, Jiangsu and Hubei) between 32° N and 40° N. Four of these provinces, Hebei, Shandong, Beijing and Tianjin, comprise a Novel Expanded Range (NER) in the northern part of the NCP that had no outbreaks during the 1950s. There were no reports of wheat midge damage in three southern provinces (Jiangxi, Zhejiang and Shanghai) below latitude 30° N in the 2010s, each of which that had recorded pest damage prior to 1960. The latitudinal median for 1950s observations was 32.74 °N (IQR 32.02 – 33.60 ° N), compared with 35.81 °N (IQR 33.81 – 37.95 °N) for the 2010s. The median latitude of midge outbreaks was 3.21 ± 0.02° SE further north in the 2010s than in the 1950s, equivalent to a range shift of 58.8 ± 0.4 km per decade (bootstrap mean of difference between median outbreak latitude in 2010s vs. 1950s). First observations of wheat midge larvae in soils at 37 sites across the NER (Supplementary Fig. S2) revealed that the insect spread northwards at a rate of 0.083 ± 0.021° N y^−1^, or 91.3 ± 23.3 km per decade since the resurgence of wheat midge in 1985 (Fig. 3). These rates of migration closely match climate velocities for the region since 1960 (*17*), and are within the observed range for dipteran crop pests in general (*1*).

**Figure 3.**
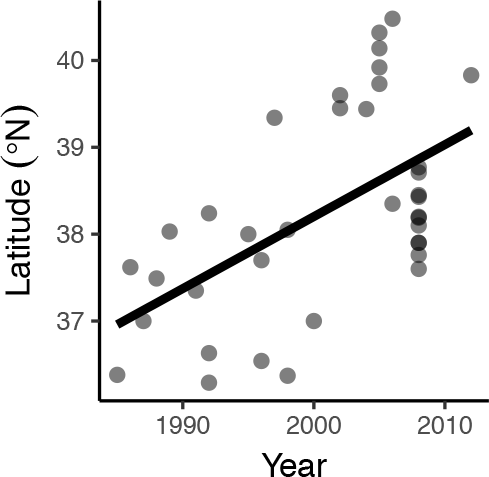
Wheat midge first arrival across the NER. Latitude of the first year in which wheat midge larvae were detected in soil in the NER, 1985-2014. Mean latitude of first observation increased by 0.083 ± 0.021 ° y^−1^ (linear regression, n = 37, r^2^ = 0.31, p < 0.001).

Poleward range shifts are expected under global warming (*2*) but we must discount any effect of changing host distribution before attributing climate change as a cause of wheat midge range shift. In the NCP, the southernmost provinces of Jiangxi and Zheijang have historically grown little wheat, declining to < 1 % of total area by the 2010s (Supplementary Fig. S3). Hence, the disappearance of wheat midge from Jiangxi and Zhejiang could be due to absence of the host. Shanghai, however, has retained 7.5 % wheat cover (down from 10.8 % in the 1950s) but did not record an outbreak in the 2010s. The central NCP provinces of Hubei, Henan, Anhui, and Jiangsu have retained moderate to high wheat cover (mean 5.9, 27.0, 14.9 and 18.2 %, respectively) with little change over time. Hebei and Shandong, which make up the majority of the NER, have relatively high and stable wheat area (12.7 and 24.3 %, respectively), while the smaller regions of Beijing and Tianjin have relatively low cover with only Tianjin showing an increase since the 1950s. Thus, the NER has always grown significant areas of wheat, and the northward range expansion of wheat midge cannot be attributed to host distribution shifts.

Wheat grain consumption by wheat midge larvae is dependent upon oviposition during wheat heading in spring (*3*). Mean spring temperatures (February, March, May temperatures from CRU gridded estimates) increased by 0.035 ± 0.007 °C y^−1^ across the NCP between 1951 and 2017, increasing more rapidly in the NER (0.045 ± 0.08 °C y^−1^) than the ODR (0.031 ± 0.06 °C y^−1^, Supplementary Fig. S4). Median spring temperatures in the 1950s for sites that experienced outbreaks in the 1950s were 8.8 °C (IQR 8.4 – 9.3 °C), whereas sites that were to experience outbreaks in the 2010s were cooler with median temperature 7.5 °C (IQR 6.6 – 8.4, Fig. 4). By the 2010s, median spring temperatures had risen to 9.3 °C (IQR 8.5 – 10.1 °C) for those sites experiencing outbreaks in the 2010s, while those which had experienced outbreaks in the 1950s had warmed to median 10.3 °C (IQR 9.8 – 10.8 °C). Thus, outbreaks are now occurring at temperatures around 0.5 °C warmer than in the 1950s. The median spring temperature during the year of arrival at each of 37 monitoring sites in the NER was 8.5 °C (IQR 7.5 – 8.9 °C).

**Figure 4.**
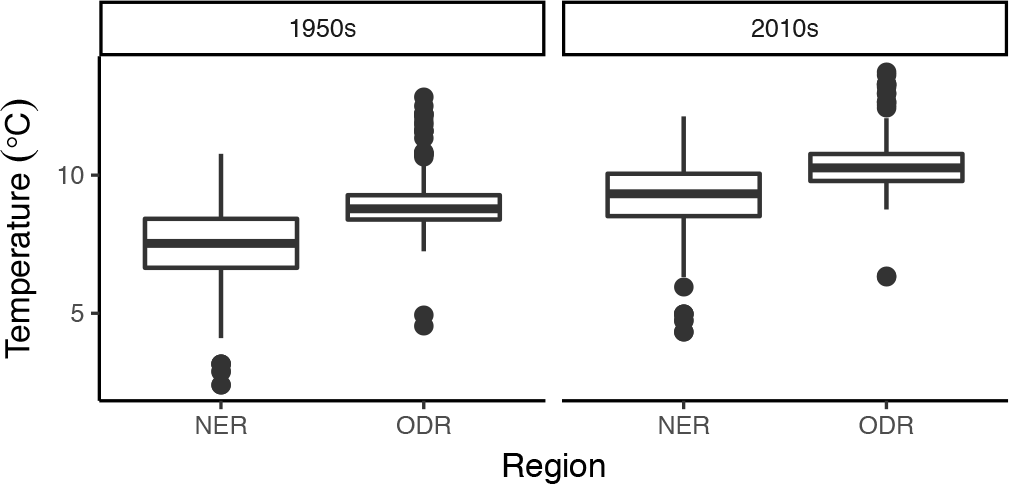
Spring temperatures of locations of wheat midge outbreaks in the ODR and NER in the 1950s and 2010s.

Wheat midge larvae consume developing wheat kernels, and adult oviposition must coincide with wheat heading, since the susceptibility of plants to wheat midge damage declines significantly after anthesis begins (*18*, *19*). Wheat and midge phenology were monitored in five sites from 30 to 39 °N in 2013, 2014 and 2016, to quantify the degree of phenological matching (Supplementary Figs. S2 & S5, Supplementary Table S1). Peak wheat heading occurred between 3^rd^ April and 6^th^ May depending on year and site, while the date by which at least half of midge adults had emerged varied between 16^th^ March and 11^th^ May. Wheat heading and adult midge emergence were delayed along increasing latitudes, by 2.93 d ± 0.37 SE per degree latitude for wheat, and 4.39 d ± 0.51 SE per degree latitude for wheat midge (estimated by linear mixed effects models with random intercepts per site). Positive mismatches (when wheat midge adults emerged before wheat heading) were observed at warmer lower latitudes, and negative mismatches (wheat heading before wheat midge emergence) at cooler, higher latitudes (Fig. 5a,b). Zero mismatch days occurred at spring temperatures of 10.1°C ± 0.4 SE (Fig. 5b). The number of larvae found in soil increased significantly with latitude (Fig. 5c, correlation r = 0.80, df = 13, p = 0.0003). The largest numbers of larvae per wheat ear were found at mismatches of −3 to 0 days, with no larvae present when phenologies were strongly mismatched (Fig. 5d). Thus, wheat midge damage is greatest at higher latitudes, where adult emergence is better synchronized with wheat heading.

**Figure 5.**
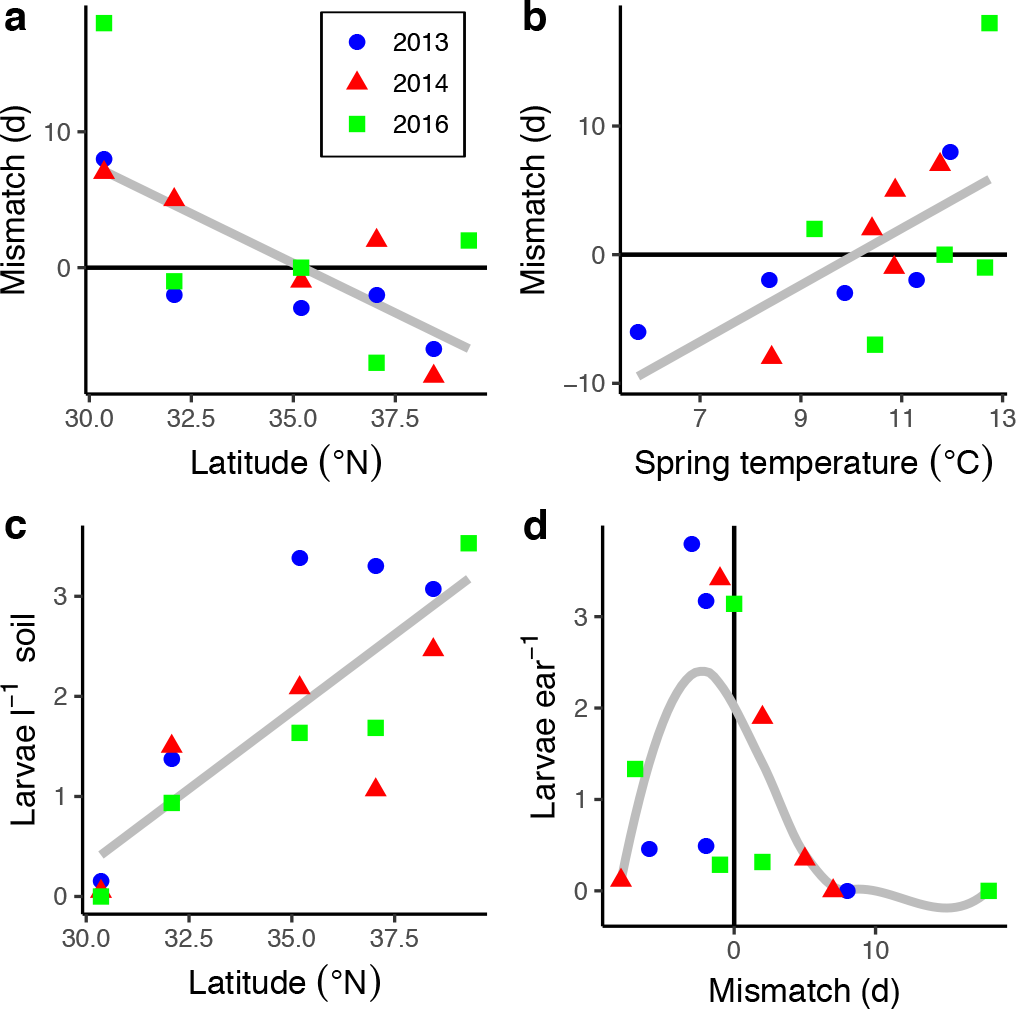
Wheat and wheat midge phenology. a) Mismatch between date of wheat midge adult emergence and date of wheat ear heading in the NCP vs. latitude, in 2013 (blue circle), 2014 (red triangle) and 2016 (green square), with line of best fit (grey). Positive mismatch indicates peak wheat midge emergence occurs before peak wheat heading, negative values that peak wheat midge emergence occurs after peak wheat heading. b) Mismatch vs. spring temperature. c) Larvae per litre of soil vs. latitude. d) Mean number of wheat midge larvae per wheat ear vs. mismatch days.

We estimated the mismatch between wheat heading and adult midge emergence across the NCP in the 1950s and 2010s, from a linear regression of mismatch days on spring temperature at the five observation sites (Fig. 5b). We applied the regression equation (mismatch days = −20.4 + 2.07 × spring temperature) to spring temperatures derived from monthly CRU temperature data (Fig. 6). The estimated mismatch days correlate with the observed distribution of midge outbreaks in the 1950s and 2010s (Fig. 1). Slightly negative mismatches, which are associated with the greatest infestations (Fig. 5d), occur in the ODR provinces of Henan, Anhui and Jiangsu during the 1950s (Fig. 6a). By the 2010s, areas of zero to slightly negative mismatches had shifted northward into the NER provinces of Hebei and Shandong (Fig. 6b). Temperature-driven phenological patterns detected in recent observational studies therefore strongly predict the locations of historical outbreaks.

**Fig. 6.**
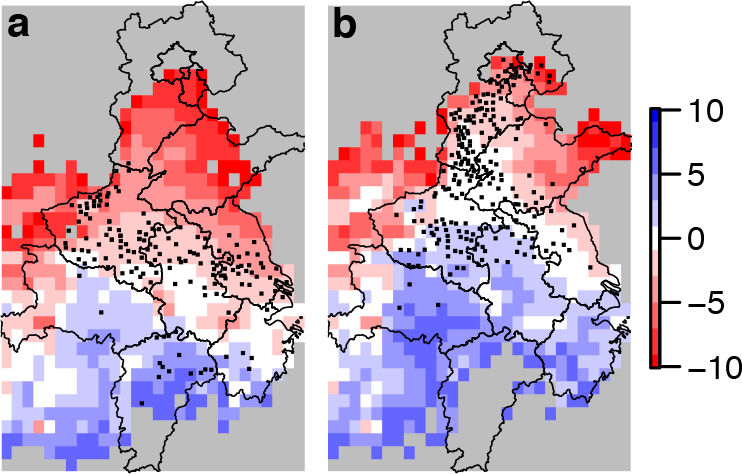
Estimated phenological mismatch between wheat heading and adult midge emergence (days) in a) the 1950s and b) 2010s. Mismatch days were estimated from mean spring temperatures (monthly CRU data) by linear regression of observational data (see text for details). Grey regions lie outside the spring temperature range in the observational study (−5 to +13 °C).

Emergence of wheat midge adults has been modelled using thermal time (cumulative degree days above a threshold temperature)(*8*, *20*) or via temperature thresholds (*9*, *21*). At our six phenology monitoring sites, adult wheat midge emergence peaked on days with a median temperature of 17.7 °C (IQR 16.6 – 19.1 °C), wheat heading peak had a much wider temperature range (median 17.7 °C, IQR 14.1 – 20.5 °C, Supplementary Fig. S6). Conversely, wheat heading occurred over a relatively small range of degree days above 5°C (median 438.5, IQR 398.5 – 460.5), compared with adult wheat midge emergence (median 447.8, IQR 391.4 – 481.9). Hence, a temperature threshold may be a more reliable predictor than temperature accumulation for adult wheat midge emergence (*9*). While some studies have suggested precipitation events as triggers for midge emergence (*9*, *10*), we found no suggestion of a relationship between rainfall and emergence (Fig. S5).

The northward range shift of wheat midge has had significant effects on food production and pest control expenditure. National Agro-tech Extension and Service Centre of China (NATESC) records show that 19.07 million ha of winter wheat was treated with insecticides in an attempt to minimize wheat midge damage in the NERC between 1985 and 2016 (Fig. 2). The cost of these insecticides (approximately US $15 per ha) means that climatic warming has caused an estimated economic loss of around $286.05 million to Chinese wheat producers during this period. Furthermore, increased insecticide usage has only been partially successful, with around 0.95 million metric tons of wheat still being lost to wheat midge damage from 1985 to 2016 (Fig. 2). Our results show that climate warming caused a northward shift in wheat midge outbreaks in the NCP. This geographical shift has resulted in the peak date of midge adult emergence coinciding more closely with winter wheat ear heading dates at higher latitudes. Wheat midge is also anticipated to expand its range in the Northern Great Plains of North America as temperatures increase, which is likely to affect spring wheat harvests over the next 40 years (*16*). There is growing interest in the use of parasitoids for biological control (*22*), acknowledging the importance of phenological matching between the parasitoid and the midge to maximize efficacy (*10*, *23*). However, little is known of the responses of these natural enemies to climate change, nor whether biocontrol could surplant chemical controls in future.

Over 80 arthropods, pathogenic microorganisms and nematode taxa could potentially cause damage to wheat species in China (*24*). The range of many of these other pest species has also expanded into northern winter wheat production areas in recent decades, with heavy pesticide use required for their control. For example, the fungal pathogen Fusarium head blight (*Fusarium graminearum*) was first detected in Henan province in 2012 (*25*). Despite the importance of these pests and diseases, their impacts, monitoring and management have often been ignored in considerations of yield improvements for wheat in China (*26*, *27*). The rapid invasion of major wheat growing regions in the NER highlights the urgency with which climate change impacts on agricultural pests and pathogens must be addressed.

## METHODS

The study spans the North China Plain (NCP), a low-lying plain ranging from 111° E to 123° E and 28° N to 40° N, and covering a total area of around 400,000 km^2^ (Fig. 1, Supplementary Fig. S1). The NCP comprises eleven provinces, of which seven (Henan, Anhui, Jiangsu, Hubei, Jiangxi, Zhejiang and Shanghai) make up our Original Distribution Range (ODR) for wheat midge, and four (Hebei, Shandong, Beijing and Tianjin) make up the Novel Expanded Range (NER).

We obtained agricultural and meteorological data for the NCP from a number of sources. Data for annual wheat production and harvested areas of each province were obtained from the crop database of the Ministry of Agricultural and Rural Affairs of the People’s Republic of China (http://202.127.42.157/moazzys/nongqing.aspx). Gridded wheat production area estimates at 10km spatial resolution for the year 2005 (used in Fig. 1) were obtained from MAPSPAM (http://mapspam.info). Monthly mean temperature data at 0.5° spatial resolution from January 1951 to December 2017 were obtained from the CRU TS v. 4.02 reanalysis (https://crudata.uea.ac.uk/cru/data/hrg/cru_ts_4.02/). Spring was defined as the months February, March and April (1 February to 30 April). For specific research sites, temperature and precipitation data were obtained from the China National Meteorological Administration (http://eng.nmc.cn/).

Analyses of wheat midge distribution and phenology were based on three datasets: an NCP-wide compilation of wheat midge outbreaks in the 1950s and 2010s; measurements of wheat midge soil larval densities, larvae per wheat ear and wheat grain loss at 37 experimental stations from 1985 onwards; and measurements of soil larval densities, adult emergence phenology, and wheat phenology at five locations across a latitudinal gradient in 2013, 2014 and 2016.

Records of location, earliest dates of 135 counties recorded damage of larvae feeding on wheat kernel in the 1950s were obtained from the Institute of Plant Protection and Soil Science, Hubei Academy of Agricultural Sciences (formerly known as the Central South Institute of Agricultural Sciences), Institute of Plant Protection, Jiangsu Academy of Agricultural Sciences (formerly known as the East China Institute of Agricultural Sciences), and from the literature (*14*, *15*, *28*). These records originate from seven provinces (Henan, Anhui, Jiangsu, Hubei, Jiangxi, Zhejiang and Shanghai). Records of 225 counties on wheat midge incidence since 2010 were obtained with permission from NATESC (https://www.natesc.org.cn/).

From 1985 to 2014, soil larval densities were sampled in 37 wheat experimental stations of the China Agricultural Research System (CARS), Plant Protection Plant and Plant Quarantine Station in Henan province, and Beijing City, giving coverage across the NER between 36° N and 40° N (Fig. S7). Soil samples (10×10×20 cm, length width and depth) were taken after annual harvesting of wheat in September. This involved diagonal sampling of 5 points in a field, with 10 randomly-selected fields per county (People’s Republic of China National Agricultural Standard NY/T616-2002, 2003). At the same sites, the fraction of wheat yield loss at each wheat grain filling stage was estimated from the number of larvae per ear (People’s Republic of China National Agricultural Standard GB/T 24501-2009). Measurements were carried out in more than 10 representative wheat fields in each of the 37 sites used for soil larval density measurements. Diagonal 5-point sampling was carried out in each field, with 10 ears of wheat collected per sampling point, and each ear checked for number of larvae and the number of grains. Percentage wheat yield losses L = 100×W/GC where W is the total number of larvae per ear, G is the total number of grains per ear, and C is the number of larvae required to consume a grain (C = 4).

Dates for wheat heading and flowering periods were recorded from March, with plots covered with 100-mesh gauze and the number of adult insect eclosions recorded daily. Close synchronization between the heading stages of winter wheat and the breeding cycle of wheat midge adults would lead to serious losses in wheat yields. Therefore, synchronization of the wheat heading period and the peak of adult insect emergence were determined for six different geographical sites along a latitudinal gradient: Wuhan (30.36 °N), Xinyang (32.08 °N), Xinxiang (35.19 °N), Xingtai (37.04 °N), Baoding (38.44 °N) and Langfang (39.30 °N) (Supplementary Fig. S5, S7, Supplementary Table S1). Boading was monitored in 2013 and 2014, after which monitoring moved to Langfang in 2016. Three main winter wheat varieties (Bima 1, Xiaoyan 4 and Zhoumai 18) were used for wheat production in these regions, with wheat damage levels determined by monitoring the number of larvae present per 100 ears in the different regions in 2013, 2014 and 2016.

## Data availability

Locations of all auxiliary datasets are listed in the Methods. All experimental datasets will be made available on DRYAD (https://datadryad.org) upon manuscript publication.

## Acknowledgements

YW was supported by the China Agricultural Research System (CARS-03) and Natural Science Foundation of Henan (182300410003). DB was funded by the Agri-tech Cornwall and Isles of Scilly programme, supported by the European Regional Development Fund, Cornwall Council, the Council for the Isles of Scilly, the Cornwall College Group, the Universities of Exeter and Plymouth and Rothamsted Research.

## Author contributions

YW, GZ, HZ, SX, GZ, DY J F, ZZ, and DY collected the data. TL, JH, HL, YJ, YJ, JD, CL, QW, DJ YZ, and JC performed the experiments. YW, ZG, DPB, JM & XL analysed the data. YW, DPB, ZG, JM and XL wrote the paper.

## References

1. D. P. Bebber, M. A. T. Ramotowski, S. J. Gurr, Crop pests and pathogens move polewards in a warming world. Nature Clim. Change. 3, 985–988 (2013).

2. D. P. Bebber, Range-Expanding Pests and Pathogens in a Warming World. Annu. Rev. Phytopathol. 53, 335–356 (2015).

3. Y. Duan et al., Genetic Diversity and Population Structure of Sitodiplosis mosellana in Northern China. PLOS ONE. 8, e78415 (2013).

4. Y. F. Li, D. J. Cai, Z. J. Shan, Z. L. Zhu, Gridded Usage Inventories of Technical Hexachlorocyclohexane and Lindane for China with 1/6° Latitude by 1/4° Longitude Resolution. Arch. Environ. Contam. Toxicol. 41, 261–266 (2001).

5. IPCC, Climate Change 2014: Impacts, Adaptation, and Vulnerability. Part A: Global and Sectoral Aspects. Contribution of Working Group II to the Fifth Assessment Report of the Intergovernmental Panel on Climate Change (Cambridge University Press, Cambridge, UK, 2014).

6. M. C. Fisher et al., Emerging fungal threats to animal, plant and ecosystem health. Nature. 484, 186–194 (2012).

7. W. Cheng, Z. Long, Y. Zhang, T. Liang, K. Zhu-Salzman, Effects of temperature, soil moisture and photoperiod on diapause termination and post-diapause development of the wheat blossom midge, Sitodiplosis mosellana (Géhin) (Diptera: Cecidomyiidae). Journal of Insect Physiology. 103, 78–85 (2017).

8. R. H. Elliott, L. Mann, O. Olfert, Calendar and degree-day requirements for emergence of adult wheat midge, Sitodiplosis mosellana (Géhin) (Diptera: Cecidomyiidae) in Saskatchewan, Canada. Crop Protection. 28, 588–594 (2009).

9. G. Jacquemin, S. Chavalle, M. De Proft, Forecasting the emergence of the adult orange wheat blossom midge, Sitodiplosis mosellana (Géhin) (Diptera: Cecidomyiidae) in Belgium. Crop Protection. 58, 6–13 (2014).

10. S. Chavalle, P. N. Buhl, F. Censier, M. De Proft, Comparative emergence phenology of the orange wheat blossom midge, Sitodiplosis mosellana (Géhin) (Diptera: Cecidomyiidae) and its parasitoids (Hymenoptera: Pteromalidae and Platygastridae) under controlled conditions. Crop Protection. 76, 114–120 (2015).

11. O. O. Olfert, M. K. Mukerji, J. F. Doane, Relationship between infestation levels and yield loss caused by wheat midge, Sitodiplosis Mosellana (Géhin) (Diptera: Cecidomyiidae), in spring wheat in Saskatchewan. The Canadian Entomologist. 117, 593–598 (1985).

12. O. Olfert, R. H. Elliott, S. Hartley, Non-native insects in agriculture: strategies to manage the economic and environmental impact of wheat midge, Sitodiplosis mosellana, in Saskatchewan. Biol Invasions. 11, 127–133 (2009).

13. J. Miao et al., Long-Distance Wind-Borne Dispersal of Sitodiplosis mosellana Géhin (Diptera:Cecidomyiidae) in Northern China. J Insect Behav. 26, 120–129 (2013).

14. X. Zeng, Wheat Midge (Agricultural Press, Beijing, 1965).

15. P. L. Yang, Research and control on Wheat Midge (Science Press, Beijing, 1959).

16. O. Olfert, R. M. Weiss, R. H. Elliott, Bioclimatic approach to assessing the potential impact of climate change on wheat midge (Diptera: Cecidomyiidae) in North America. The Canadian Entomologist. 148, 52–67 (2016).

17. M. T. Burrows et al., The pace of shifting climate in marine and terrestrial ecosystems. Science. 334, 652–655 (2011).

18. Y. Wu et al., The synchronization of ear emerging stages of winter wheat with occurrent periods of the orange wheat blossom midge, Sitodiplosismosellana (Gehin) (Diptera: Cecidomyiidae) adults and its damaged level. Acta Ecologica Sinica. 35, 3548–3554 (2015).

19. R. H. Elliott, L. W. Mann, Susceptibility of red spring wheat, Triticum aestivum L. cv. Katepwa, during heading and anthesis to damage by wheat midge, Sitodiplosis mosellana (Géhin) (Diptera: Cecidomyiidae). The Canadian Entomologist. 128, 367–375 (1996).

20. J. F. Doane, O. Olfert, Seasonal development of wheat midge, Sitodiplosis mosellana (Géhin) (Diptera: Cecidomyiidae), in Saskatchewan, Canada. Crop Protection. 27, 951–958 (2008).

21. J. N. Oakley et al., Prediction of orange wheat blossom midge activity and risk of damage. Crop Protection. 17, 145–149 (1998).

22. S. Chavalle, P. N. Buhl, G. San Martin y Gomez, M. De Proft, Parasitism rates and parasitoid complexes of the wheat midges, Sitodiplosis mosellana, Contarinia tritici and Haplodiplosis marginata. BioControl. 63, 641–653 (2018).

23. R. H. Elliott, L. Mann, O. Olfert, Calendar and degree-day requirements for emergence of adult Macroglenes penetrans (Kirby), an egg-larval parasitoid of wheat midge, Sitodiplosis mosellana (Géhin). Crop Protection. 30, 405–411 (2011).

24. Y. Y. Guo, Crop diseases and insect pests (China Agricultural Press, Beijing, 2015).

25. F. Xu et al., First Report of Fusarium pseudograminearum from Wheat Heads with Fusarium Head Blight in North China Plain. Plant Disease. 99, 156–156 (2014).

26. X. Qin et al., Wheat yield improvements in China: Past trends and future directions. Field Crops Research. 177, 117–124 (2015).

27. S. Sun, X. Yang, X. Lin, G. F. Sassenrath, K. Li, Climate-smart management can further improve winter wheat yield in China. Agricultural Systems. 162, 10–18 (2018).

28. C. C. Long, White midge S. mosellana in Jianxi Province. Bulletin of Agricultural Sciences. 3, 147 (1955).

