## Supplementary Material for "Phenological matching drives wheat pest range shift under climate change"

Wu et al.

**Supplementary Figures**

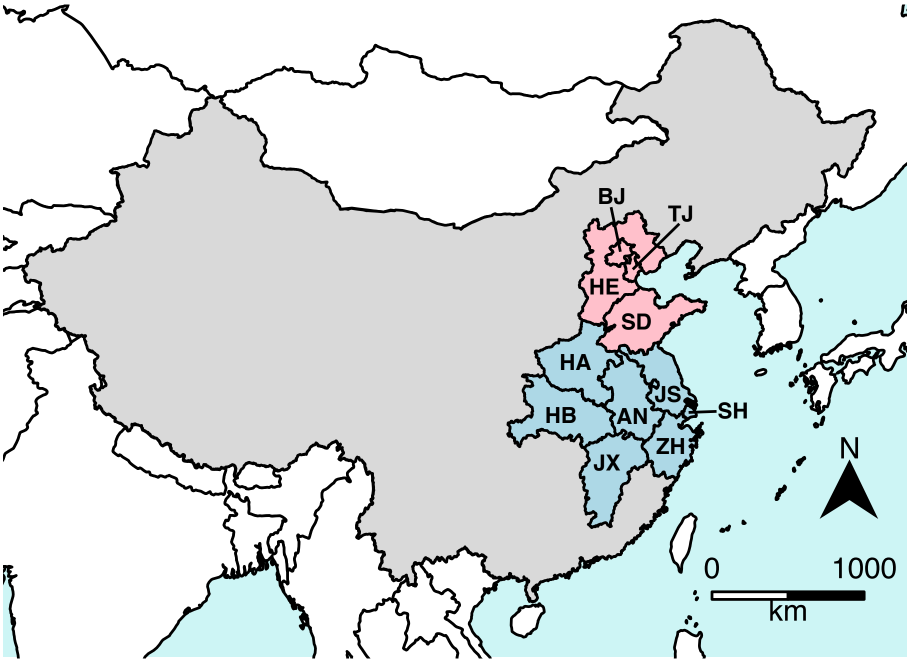


Figure S1. Location of the NCP Provinces (grey) within China. Labelled provinces and districts are Anhui (AH), Beijing (BJ), Henan (HA), Hebei (HE), Hubei (HB), Jiangsu (JS), Jiangxi (JX), Shandong (SD), Shanghai (SH), Tianjin (TJ) and Zhejiang (ZH).

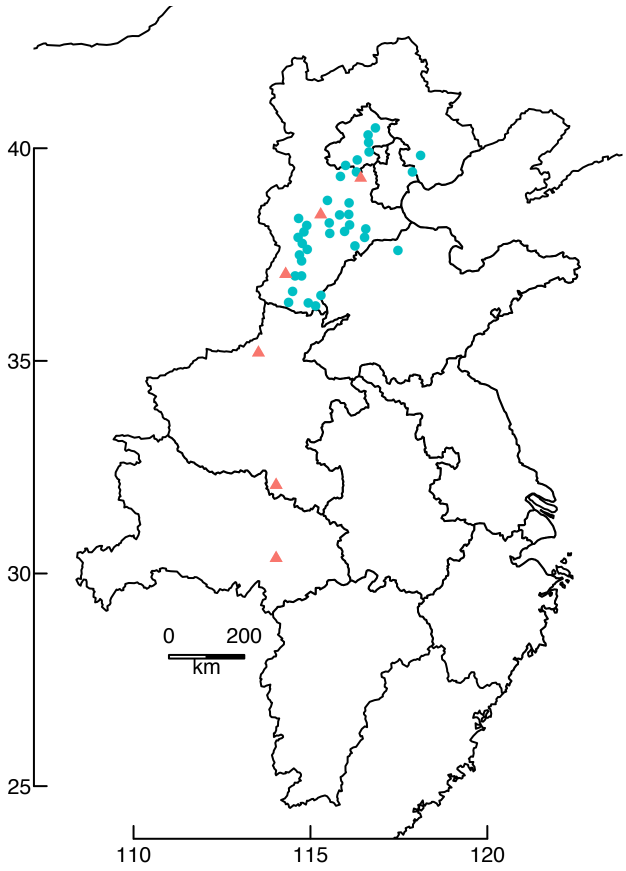


Figure S2. Locations of 37 sampling sites of wheat midge larval density (blue circles) and investigation of mismatches between wheat ear heading and wheat midge adult emergence and wheat ear damage (red triangles).

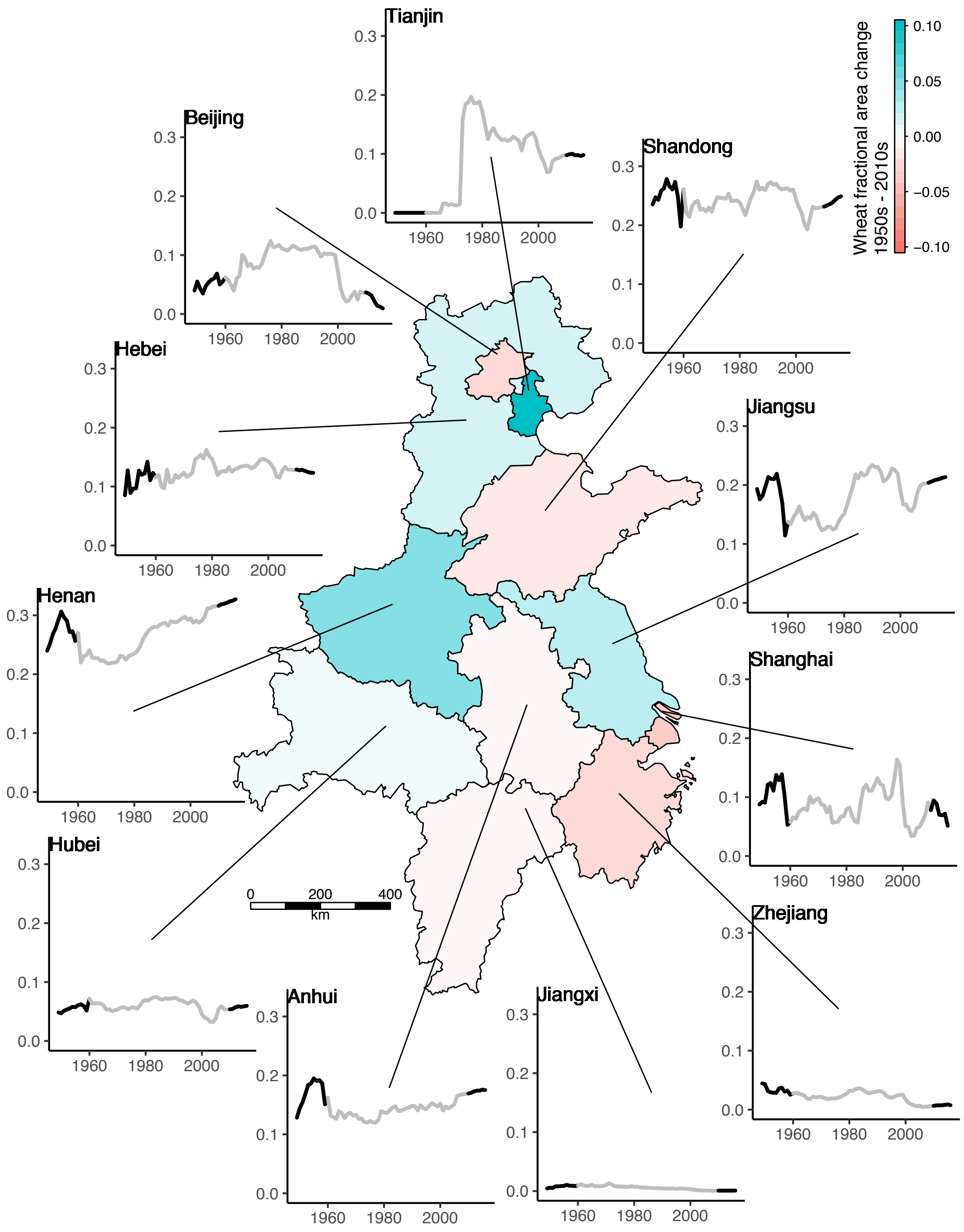


Figure S3. Change in wheat area 1949-2016 across the NCP. Graphs show fractional areal coverage by wheat production per province, with 1950s and 2010s in black, other years in grey. Map shows difference in fractional coverage between 1950s and 2010s. The most southerly Provinces of Jiangxi and Zhejiang have historically had the lowest coverage, which has declined. Central Provinces of Henan, Anhui, Jiangsu and Shandong have the highest coverage, which has varied little over time.

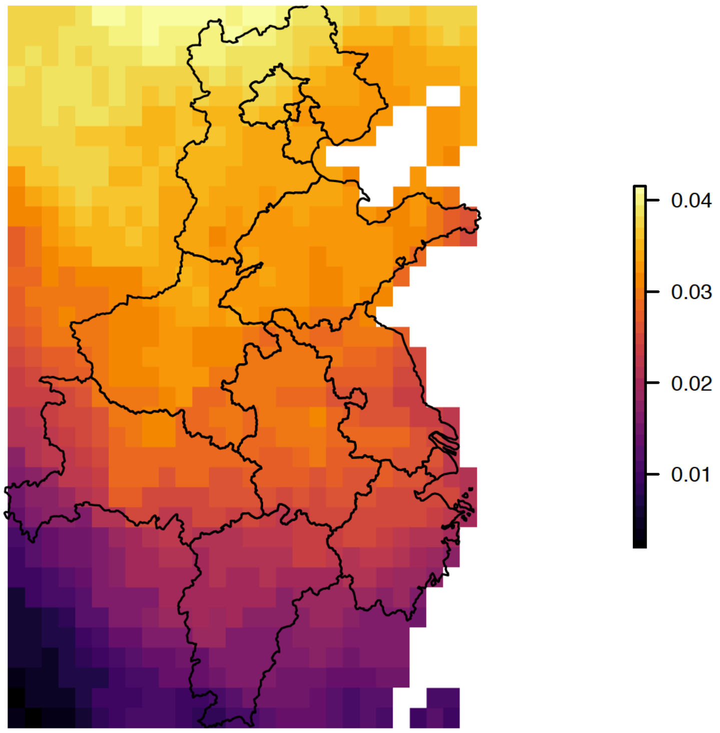


Figure S4. Spring warming trend 1951 – 2017, °C $y^{-1}$ for the NCP, data from CRU TS4.2 monthly temperature dataset. The warming trend increases with latitude.

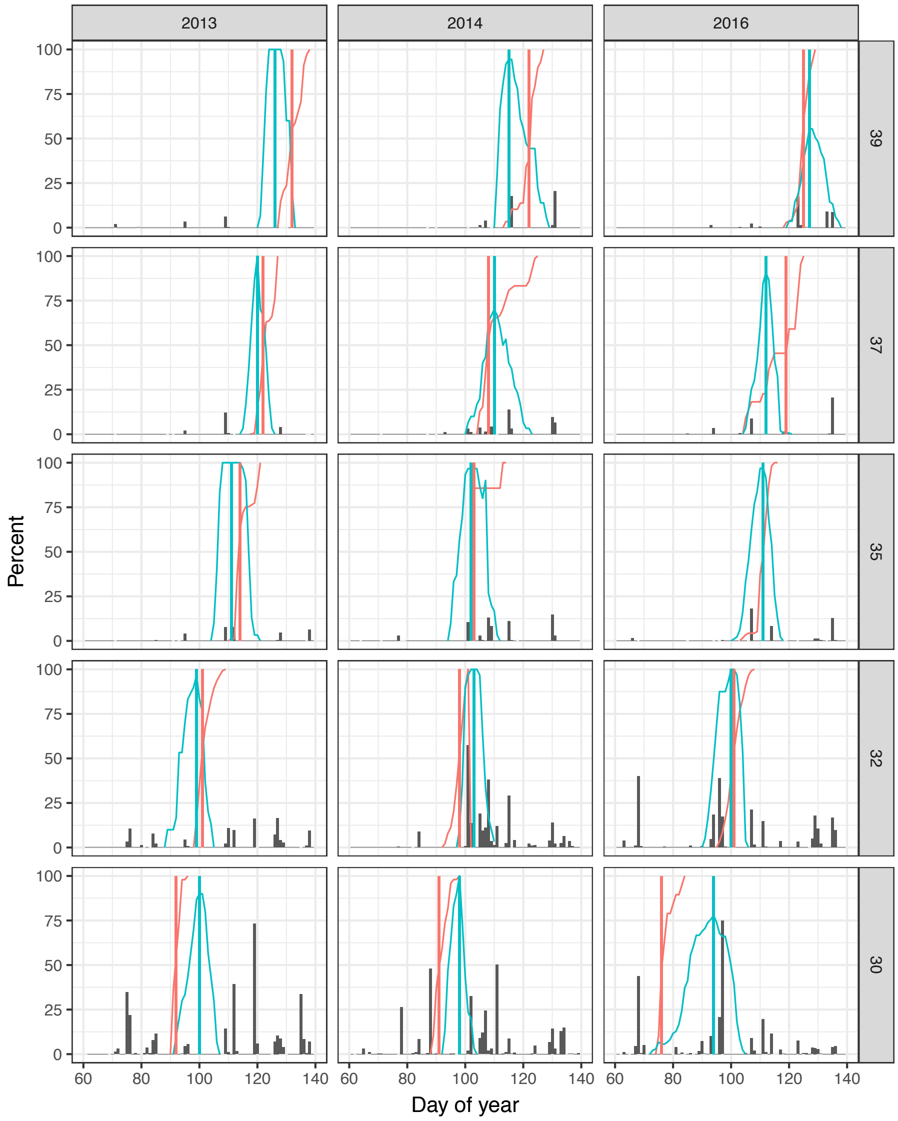


Figure S5. Wheat heading and wheat midge adult emergence by day of year at different latitudes. Blue curves show the percentage of wheat at the ear emergence stage (growth stage GS51 – GS59) in the Xiaoyan 6 wheat cultivar. Blue vertical lines show the peak stage of wheat ear emergence, designated as the day on which the percentage of wheat at the ear emergence stage is greatest, or the middle of the period of maximum emergence where there are several consecutive days. Red curves correspond to the cumulative emergence percentage for wheat midge adults. Red vertical lines give the date by which at least 50 % of adults have emerged. In general, midges emerge before wheat heading at low latitudes, and after wheat heading at high latitudes.

Columns indicate the year of sampling, and rows the latitude of the experimental area: Baoding (Hebei) at 39.0 °N, 115.5 °E for 2013 and 2014; Langfang (Hebei) at 39.5 °N, 116.6°E from 2016; Xingtai (Hebei) at 37 °N 114.5 °E; Yuangyang (Henan) at 35.0 °N, 113.7 °E; Xinyang (Henan) at 32.0 °N, 114.1 °E; Wuhan, Hubei at 30.5 °N, 114.3 °E. Locations are shown in Fig. S3.

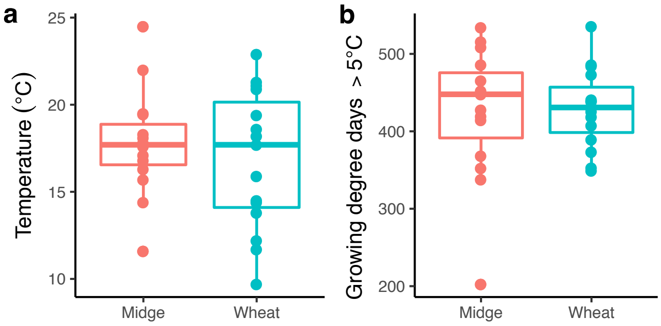

Fig. S6. a) Spring temperature and b) Growing degree days for wheat and midge phenology at five phenology monitoring sites over three years (2013, 2014, 2016). Boxes show interquartile ranges, horizontal bars show medians.

### Supplementary Tables

Table S1. Phenological sites. Code refers to the Chinese meteorological station number. Data for Langfang were unavailable so the means for nearby Baodi and Bazhou were used.

| Site | Code | Province | Latitude (°N) | Longitude (°E) | Altitude (m) |
|---|---|---|---|---|---|
| Baodi | 54525 | Tianjin | 39.44 | 117.17 | 5.1 |
| Langfang | 54515 | Hebei | 39.30 | 116.42 | 13.7 |
| Bazhou | 54518 | Hebei | 39.10 | 116.24 | 8.9 |
| Xingtai | 53798 | Hebei | 37.04 | 114.30 | 77.3 |
| Baoding | 54602 | Hebei | 38.44 | 115.29 | 16.8 |
| Xinxiang | 53986 | Henan | 35.19 | 113.53 | 73.2 |
| Xinyang | 57297 | Henan | 32.08 | 114.03 | 114.5 |
| Wuhan | 57494 | Hubei | 30.36 | 114.03 | 23.6 |
